# CRISPR-Cas13d-Mediated Targeting of a Context-Specific Essential Gene Enables Selective Elimination of Uveal Melanoma

**DOI:** 10.1101/2025.08.21.671629

**Authors:** Daniel Stauber, Lucas Sosnick, Yitong Ma, Sopida Pimcharoen, Atip Lawanprasert, Niren Murthy, David Myung, Lei Stanley Qi

## Abstract

Uveal melanoma, the most common eye cancer in adults, remains limited to surgical intervention and chemotherapy, with a dismal survival rate that has not improved in over 50 years. To address this therapeutic impasse, we systematically analyzed public gene expression, RNAi, and CRISPR knockout datasets and identified RASGRP3 as an essential gene specifically for uveal melanoma. RasGRP3 is uniquely overexpressed and essential for survival in uveal melanoma cells, but dispensable in healthy cells. RasGRP3 remains “undruggable” due to its intracellular localization and lack of targetable binding pockets. To overcome this, we developed a CRISPR-Cas13d RNA-targeting therapeutic that specifically knocks down RasGRP3 mRNA. This Cas13d-based therapeutic mediates selective uveal melanoma killing through two synergistic mechanisms: (i) direct silencing of the essential RasGRP3 transcript, and (ii) collateral RNA degradation triggered by the cleavage of overexpressed RasGRP3. When delivered via optimized lipid nanoparticles encoding Cas13d mRNA and guide RNA, this strategy eliminated >97% of uveal melanoma cells while sparing healthy cells, including retinal pigment epithelial cells. This approach outperformed conventional Cas9 and siRNA methods in potency without inducing permanent genomic alterations. Our findings establish a RNA-targeting therapeutic for uveal melanoma and a framework for Cas13d-based interventions against broad “undruggable” cancers.

## Introduction

Uveal melanoma (UM) is the most common primary intraocular malignancy in adults and the second-most common melanoma in the world, accounting for up to 5% of all melanoma cases.^1,2^ It arises from melanocytes in the iris, ciliary body, or—most commonly—the choroid (>90%). The prognosis is poor: 50% of all patients eventually develop metastatic disease, predominantly to the liver, and median survival time after metastasis is 6-12 months.^3^ Notably, the 5-year survival rate has not improved for over half a century.^4–6^

Standard-of-care treatment still begins with radiation therapy (plaque brachytherapy or proton beam irradiation), often followed by enucleation (surgical eye removal).^1,7^ Once metastasis occurs, a single FDA-approved immunotherapy, Tebentafusp, improves survival time by just ∼5 months, and only works in a minority of patients (∼33% of cases, which are HLA-A*02:01-positive),^8–10^ leaving the majority of those afflicted with virtually no options.^6,11,12^ The dismal therapeutic landscape, coupled with the devastating visual and psychological consequences of standard-of-care treatments,^7^ highlights an acute need for innovative precision therapeutics.

Oncogenic drivers of UM are well catalogued—mutations in GNAQ/GNA11 activate PLCβ-PKC signaling, triggering constitutive MAPK output—however, targeting the core proteins of this cascade (e.g., PKC, MEK) has yielded only modest clinical benefit.^13–15^ High-throughput CRISPR knockout screens (e.g., DepMap)^16–19^ now enable systematic interrogation of context-specific essential genes: genes whose loss is lethal in one cancer lineage yet dispensable in normal tissues.^18,20^ Mining ∼2,000 known fitness genes with a custom statistical pipeline, we have distilled the list to over 60 previously undrugged, context-specific targets (**Fig. 1A**). Among these, Ras Guanyl Releasing Protein 3 (*RASGRP3*) surfaced as a top hit for UM. As a guanine nucleotide exchange factor, RasGRP3 acts directly downstream of PKC to load GTP onto Ras, amplifying oncogenic MAPK signaling. Because RasGRP3 lacks obvious small-molecule pockets, resides intracellularly, and has no identified enzymatic active site, it is considered “undruggable” by conventional medicinal chemistry methods.^21,22^ Recent work using small interfering RNA (siRNA) to knock down RasGRP3 resulted in the selective killing of uveal melanoma cells, but achieved only moderate knockdown efficiency and exhibited off-target effects that would limit clinical translation.^13,23^ Nonetheless, its selective dependency profile and elevated expression in eye cancers relative to normal tissues establish RasGRP3 as an ideal proof-of-concept target for a new therapeutic modality.

**Figure 1:**
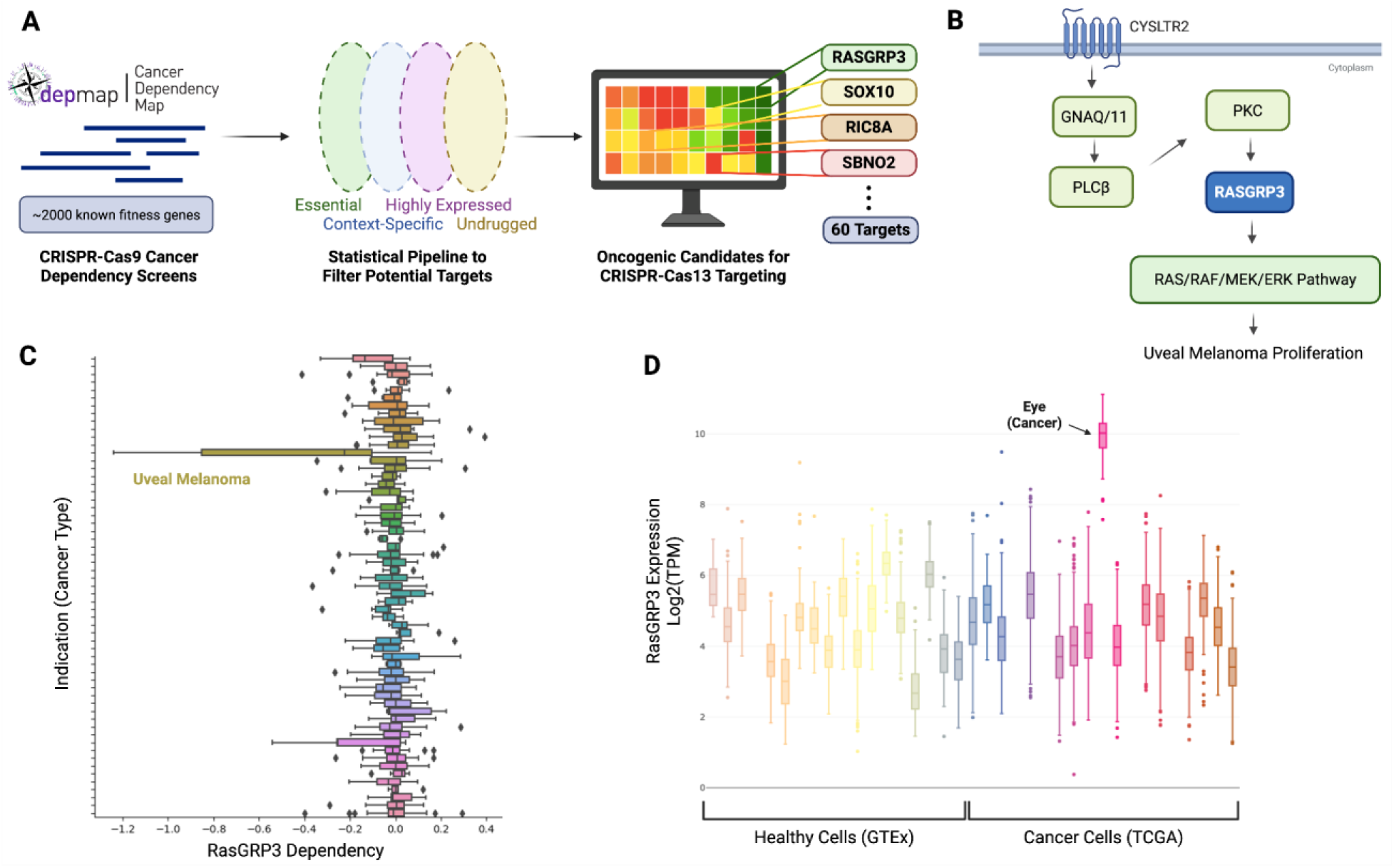
Bioinformatic screening reveals RasGRP3 as a highly-expressed context-specific essential gene for uveal melanoma. (A) Diagram of target identification pipeline using CRISPR-Cas9 cancer dependency screens. (B) Uveal melanoma proliferation pathway and the function of RasGRP3. (C) CRISPR-Cas9 knockout screen data reveals that uveal melanoma is highly dependent on RasGRP3. (D) RNA-seq data shows that eye cancer (from TCGA) exhibits several-fold higher expression of RasGRP3 than both other cancers and healthy tissues (GTEx).

Type VI CRISPR enzymes, such as Cas13d, are RNA-guided RNases that can be re-targeted with a CRISPR-associated guide RNA (gRNA) to bind and degrade complementary mRNA.^24–26^ Unlike Cas9, Cas13d does not introduce permanent edits at the DNA level; and unlike siRNA, Cas13d exhibits minimal off-target binding.^25,27,28^ Additionally, it is well-documented that Cas13 effectors can induce “collateral” or trans-cleavage of bystander transcripts following the cleavage of highly abundant primary targets.^29–31^ Although this collateral activity is often considered undesirable, we hypothesize that it can be harnessed to enhance tumor cell killing, specifically in cancer cells with high expression of the target gene—such as RasGRP3 in UM—while sparing normal cells with low or absent expression. As with many targeted therapies, inhibiting a single gene such as RasGRP3 may lead to resistance mechanisms in cancer cells. However, our Cas13 system mitigates this risk by activating a secondary cell death pathway through collateral RNA degradation, which may be critical for sustained therapeutic efficacy.

Efficient, safe, and scalable delivery of CRISPR effectors remains a central bottleneck. Adeno-associated viral vectors are limited by payload size, integration risk, and immunogenicity, whereas lentiviral or adenoviral approaches are unsuitable for transient expression.^32^ Lipid nanoparticles (LNPs)—validated clinically by mRNA vaccines like those for COVID-19^33^—can encapsulate large RNA payloads, self-assemble in a single-step microfluidic process, and be dosed repeatedly with minimal innate immune activation.^34–36^ State-of-the-art LNPs have already transported Cas9 mRNA to the liver in clinical trials, and recent mouse studies have demonstrated delivery into ocular tissue and extended this platform to Cas13d.^37–39^ Here, we hypothesize that LNP-mediated Cas13d mRNA delivery serves as a safe and efficient delivery method to UM cells.

The convergence of (i) an identified context-specific essential gene (*RASGRP3*), (ii) a programmable RNA-targeting nuclease (Cas13d) capable of transient yet potent knock-down, and (iii) a clinically-validated delivery vehicle (LNP) motivated our strategy: that LNP-mediated co-delivery of Cas13d mRNA and a RasGRP3-targeting gRNA will induce highly-specific elimination of uveal melanoma while sparing healthy tissues. The work presented here integrates large-scale cancer dependency datasets, Cas13d gRNA optimization, and LNP formulation screening to create and test a proof-of-concept Cas13d-based therapeutic for uveal melanoma. This integrated approach leads to an optimized Cas13d-LNP therapeutic that ablates >97% of UM cells *in vitro* while sparing non-UM controls. Finally, we analyzed the mechanism of cytotoxicity and confirmed the contribution of collateral RNA degradation.

Collectively, these results establish a potential first-in-class therapeutic strategy for uveal melanoma. We believe the approach demonstrated here may extend well beyond uveal melanoma and lay the foundation for broader application of RNA-targeting CRISPR systems in oncology.

## Results

### Computational Identification of RasGRP3 as a Potential Therapeutic Target for Uveal Melanoma

To identify therapeutically relevant cancer targets for an RNA-targeting approach, we first performed a comprehensive analysis of the Broad Institute’s DepMap (CRISPR-Cas9 knockout screens) comprising over 1,000 cancer cell lines spanning >30 indications (**Fig. S1A**).^40^ This analysis proceeded through four sequential filters: essentiality, context-specific lethality, high relative expression, and druggability (**Fig. 1A**, **Fig. S1B**).

First, we retained genes with a Chronos (dependency) score < −0.6 for at least one cell line, yielding greater than 2,000 genes that are essential for any number of cells—ranging from pan-cancer fitness genes to cell line-specific essential genes. Next, cell lines were grouped into their respective cancer indications. Genes with significantly higher dependency scores (in two of three independent tests, FDR-adjusted p < 0.05) were designated context-specific essential genes. These genes were then manually cross-referenced with bulk RNA-seq from TCGA^41,42^ (cancer cells) and GTEx^43^ (normal/healthy tissues). Only genes whose median expression in the target cancer exceeded both the pan-cancer and healthy-tissue medians by ≥ 2-fold (log_2_ scale) were further analyzed. Finally, genes with FDA-approved inhibitors, advanced clinical candidates, or obvious ligandable pockets (via Open Targets^44^) were removed to focus on undrugged targets. This pipeline compressed ∼17,000 assayed genes to ∼60 high-confidence, undrugged context-specific essential genes for various cancers (**Fig. 1A, Supp. Data 1**). Notably, the uveal melanoma lineage displayed the strongest enrichment for Ras guanyl releasing protein 3 (RasGRP3).

RasGRP3’s dependency profile was highly selective: across eight independent UM cell lines, the mean score was −0.76 (median: −0.25), whereas all other (>1,000) lines clustered around neutrality (**Fig. 1C**). The difference remained significant after strict multi-test correction (q < 3×10^-13^). Parallel RNA-seq analysis revealed a unique expression pattern: eye-derived tumors in TCGA (predominantly UM tumors) expressed RasGRP3 at >8-fold above every GTEx healthy tissue and every other TCGA cancer type (**Fig. 1D**).

Pathway mapping rationalized this context-specific essentiality of RasGRP3 (**Fig. 1B**). Mutations in GNAQ/GNA11 are the most common oncogenic mutations linked to UM. These mutations result in activation of the RAS/RAF/MEK/ERK cascade and further lead to UM proliferation. RasGRP3 serves as the primary guanine-nucleotide exchange factor (GEF) directly linking the upstream GNAQ/GNA11 mutations to the downstream RAS pathway activation.^13,45^ Thus, RasGRP3 is frequently and necessarily upregulated in these GNAQ/GNA11 mutant UM cells to induce proliferation. Moreover, loss of RasGRP3 therefore cripples proliferation uniquely in UM, whereas other tissues leverage alternative GEFs to maintain RAS/RAF/MEK/ERK signaling and proliferation.

*In-silico* tractability (via Open Targets^46^) classified RasGRP3 as undruggable: no known drugs or bioactive small molecules, limited structural coverage (average pLDDT = 69.87), and an intracellular localization that precludes most antibody modalities (**Fig. S1B**). However, CRISPR-based technologies can overcome these barriers by operating at the RNA level, making RasGRP3 an ideal proof-of-concept target for a Cas13 therapeutic.

### Screening for an Optimal Cas13d Guide RNA to Target RasGRP3

To transiently knock down RasGRP3 mRNA, we employed CRISPR-Cas13d, an RNA-guided ribonuclease that uses a complementary guide RNA (gRNA) to bind and cleave its target transcript.^25^ A unique feature of Cas13d is its collateral cleavage activity: once the target transcript is cleaved, Cas13d non-specifically degrades bystander RNAs (**Fig. 2A**).^29,30^ This collateral effect is dependent on the abundance of the target transcript; greater target cleavage leads to increased bystander RNA degradation, which is associated with enhanced cell killing. We hypothesized that this collateral activity could amplify tumor cell killing following essential transcripts knockdown.

**Figure 2:**
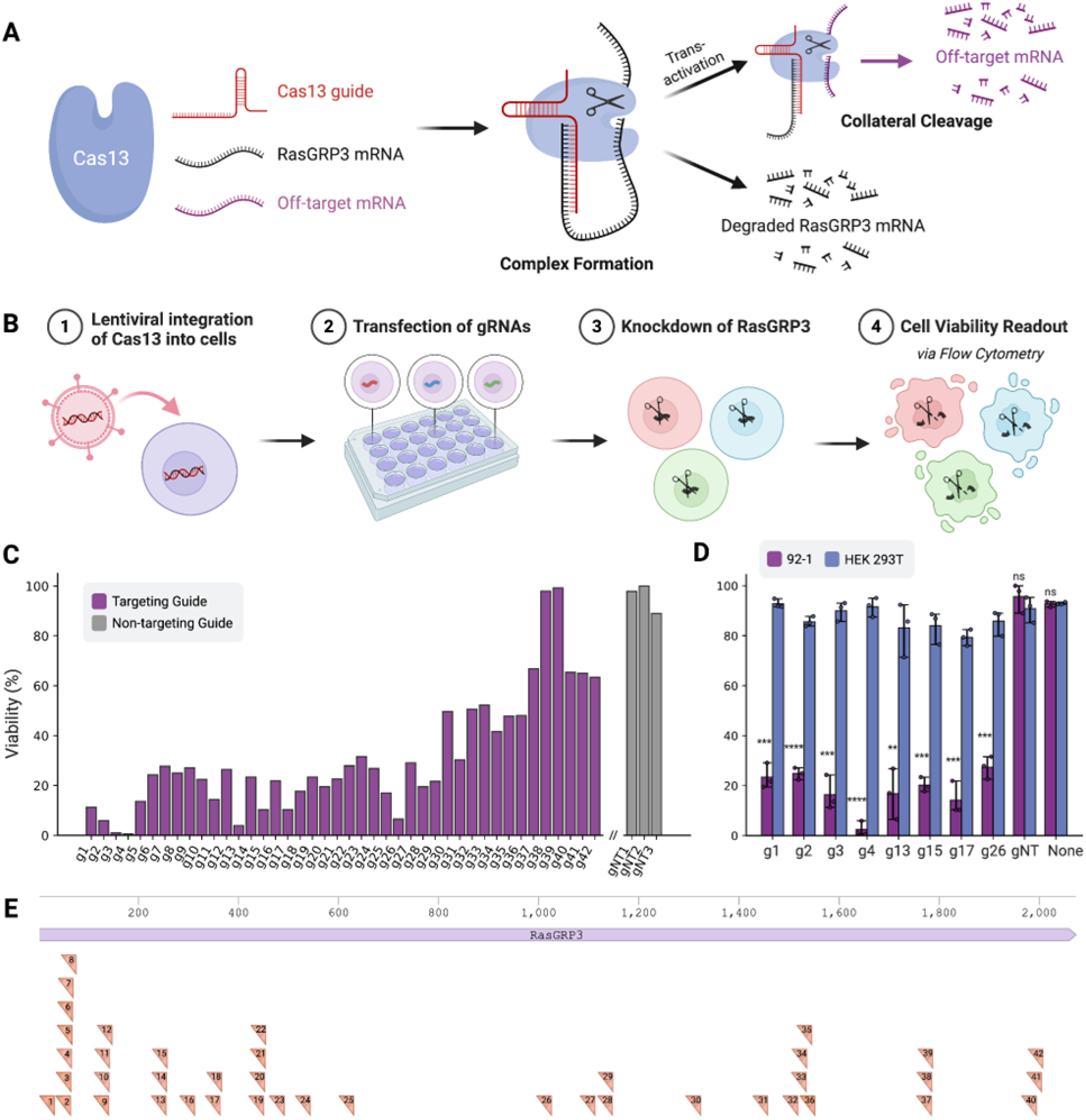
Targeting RasGRP3 using CRISPR-Cas13d yields potent and selective killing of uveal melanoma cells. (A) Diagram of Cas13’s mechanism of action: once the gRNA binds to the target mRNA, it will degrade the target mRNA and surrounding off-target mRNA. (B) Schematic of methods used in (C) and (D). Cas13d is lentivirally transduced into cells, followed by transfection of individual guides to induce knockdown of RasGRP3. Cell viability is quantified 48 hours after transfection. (C) Viability assay in 92-1 (UM) cells to screen the efficacy of 42 different gRNAs targeting RasGRP3, along with three non-targeting gRNAs. (D) Viability assay in both 92-1 (UM) and HEK 293T (non-UM) cells to examine gRNA specificity using the eight best gRNAs from (C), along with a non-targeting gRNA and untransfected control. Each data point represents one biological replicate. (E) RasGRP3 gene map with each targeting gRNA screened in (C) annotated based on the location it targets.

To pinpoint optimal gRNAs against RasGRP3, we designed 42 guides spanning the RasGRP3 coding sequence using two independent algorithms (e.g., Deep Learning-Powered Cas13d Guide Design^47,48^ and Cas13design^49^) and manual curation (**Supp. Table S1**). We used 92-1 and MP-46 cells, two lines derived from primary human uveal melanoma (accession CVCL_8607 and CVCL_4D13, respectively—Cellosaurus database), for our initial guide screening. The screening strategy included first integrating a constitutive Cas13d expression plasmid into 92-1 cells via lentiviral transduction. The transduction efficiency of Cas13d was quantified prior to screening via flow cytometry, demonstrating high expression in three different cell lines (HEK-293T: 99.1%; 92-1: 96.7%; MP-46: 84.2%) (**Fig. S2**). Given the slightly lower efficiency in MP-46 cells, subsequent screening was performed using 92-1 cells. Individual gRNAs targeting RasGRP3 were transfected into Cas13d-transduced 92-1 cells, leading to the subsequent knockdown of RasGRP3 (**Fig. 2B**). Viability varied from 0.61% (g4) to 97.93% (g37) among the recipient cells transfected with targeting gRNAs, while cells with non-targeting gRNAs all showed >85% viability (**Fig. 2C**). This established a wide dynamic range and showed that Cas13d-mediated knockdown of RasGRP3 mRNA can effectively kill uveal melanoma cells.

To test the specificity of killing, the top eight RasGRP3-targeting gRNAs (determined via lowest cell viabilities) were then retested in triplicate across 92-1 and HEK293T cells (**Fig. 2D**). Out of these, “g4” showed the best UM killing (viability at 2.50 ± 2.95%) in 92-1 cells. The data also showed that nearly all the guides had minimal effect on HEK293T cells, which remained at >80% viability among all the conditions tested, demonstrating that RasGRP3-mediated knockdown via Cas13d can selectively kill uveal melanoma cells. The 42 RasGRP3-targeting gRNAs are depicted tiled along the RasGRP3 coding sequence; the top-performing guide (g4) targets the beginning (nucleotides 38-67) of the RasGRP3 coding sequence (**Fig. 2E**). Based on its combination of potency and selectivity, g4 was chosen as the lead guide for all subsequent experiments.

### LNP Enables Efficient Delivery of Cas13d mRNA to Various Uveal Melanoma Cell Lines

Given the low immunogenicity of lipid nanoparticles (LNPs) and the liver tropism of UM metastases,^33,36^ we sought to develop an LNP-based delivery vehicle for Cas13d mRNA and RasGRP3-targeting gRNA (g4). We first optimized LNP composition for maximal mRNA transfection into UM cells, testing 30 LNP formulations that have been FDA-approved or clinically tested (**Supp. Table S2**). The low toxicity of RasGRP3 knockdown in cells other than UM cells allowed us to focus our LNP screening on high efficiency, instead of cell type or tissue specificity.

GFP mRNA alone was packaged into distinct LNP formulations—each varying in ionizable lipid, phospholipid, cholesterol, and PEG-lipid ratios—and transfected into two representative UM lines (MP-46 and 92-1) in parallel (**Fig. 3A**). Transfection efficiencies were quantified via flow cytometry, and Formulation #7 (3060i10) emerged as the clear lead—achieving around 90% GFP+ cells in both lines—while several LNP formulations (e.g., #12) yielded < 5% GFP+ cells (**Fig. 3B**). We further confirmed LNP 3060i10 delivers GFP to a list of UM (92-1, MP-46, and MP-41) and non-UM (HEK-293T, HeLa, and ARPE-19) cell lines (used below to test our therapeutic) with high efficiency and minimal toxicity (**Fig. S3**).

**Figure 3:**
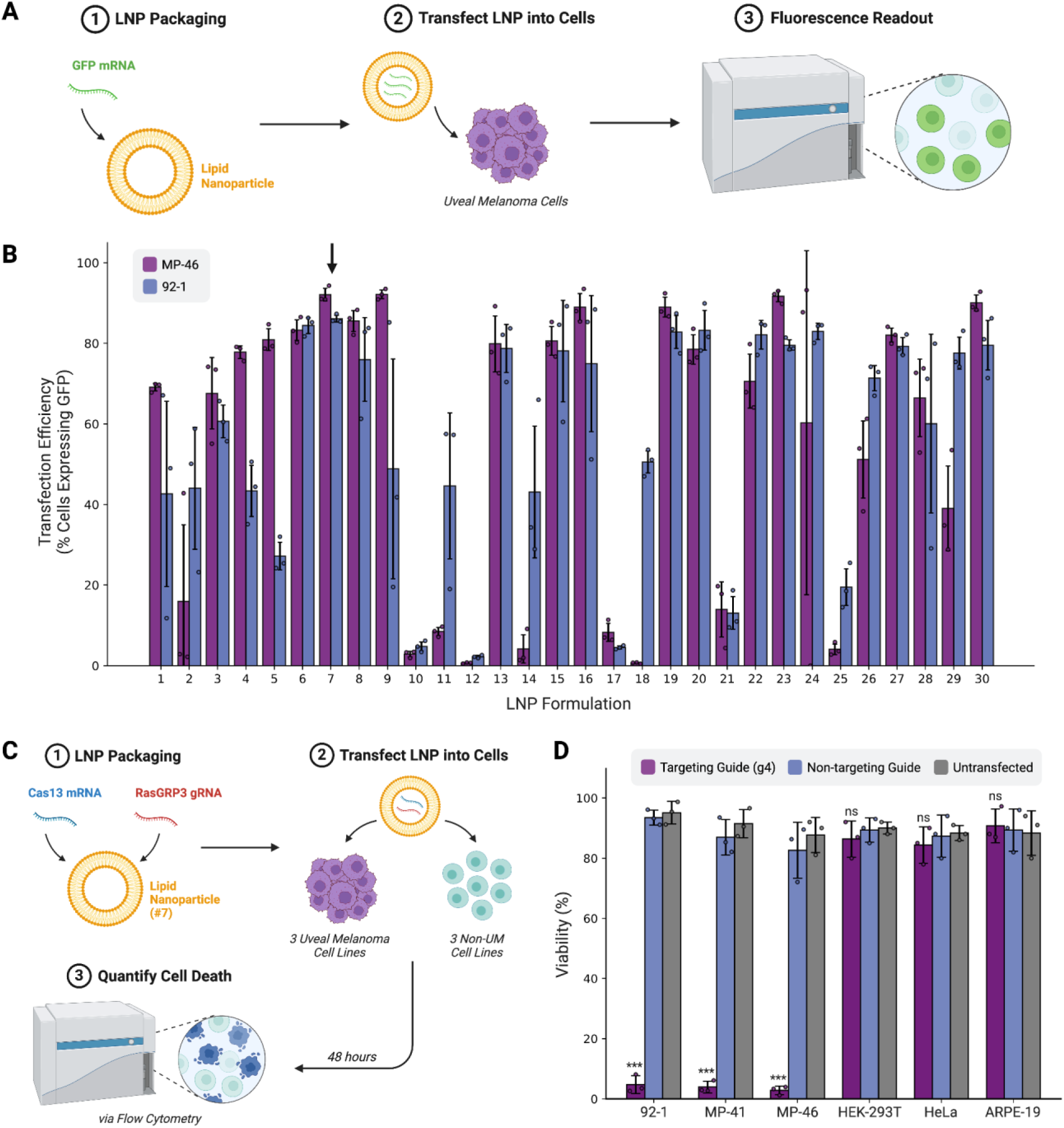
Cas13d mRNA and RasGRP3-targeting gRNA delivered via a therapeutically relevant LNP induce UM-specific cell death. (A) Schematic of methods for (B). GFP mRNA is packaged into each LNP and transfected into uveal melanoma cells to examine delivery efficiency via fluorescence. (B) Transfection efficiency of 30 different LNP formulations in two uveal melanoma cell lines (MP-46, 92-1). The arrow is pointing to Formulation #7 (3060i10), which is used in subsequent experiments. Each data point represents one biological replicate. (C) Schematic of methods for (D). The therapeutic is formed by encapsulating Cas13d mRNA and RasGRP3-targeting gRNA in an LNP. The LNP is transfected into three uveal melanoma cell lines and three non-uveal melanoma cell lines, and then cell death is measured via flow cytometry after 48 hours. (D) In-vitro viability assay in three uveal melanoma cell lines (92-1, MP-41, MP-46) and three non-UM cell lines (HEK-293T, HeLa, ARPE-19). Each data point represents one biological replicate.

Next, Cas13d mRNA and g4 gRNA were co-packaged into LNP 3060i10 (forming the final “Cas13d-LNP formulation”) and transfected into three UM cell lines (92-1, MP-46, and MP-41) alongside three non-UM controls (HEK293T, HeLa, and ARPE-19) (**Fig. 3C**). Notably, ARPE-19 is an epithelial cell line resembling uveal melanoma cells before malignancy (i.e., healthy eye cells). Remarkably, all three UM cell lines exhibited < 5% viability (92-1: 4.73 ± 2.95%; MP-41: 3.90 ± 1.91%; MP-46: 2.80 ± 1.42%), representing approximately 20-fold more killing compared to non-targeting controls (**Fig. 3D**). In contrast, non-UM lines showed minimal cytotoxicity, with viabilities (HEK293T: 86.33 ± 6.03%; HeLa: 84.34 ± 5.13%; ARPE-19: 90.73 ± 5.62%) similar to the untransfected controls. This suggests two important conclusions: (i) the Cas13d-LNP therapeutic achieves nearly complete elimination of UM cells while sparing non-UM cells, and (ii) the LNP alone has a negligible effect on viability.

### Cas13d-Mediated Knockdown of RasGRP3 Engages Collateral Cleavage to Outperform Cas9 and siRNA in Tumor Killing

To dissect the mechanisms underlying Cas13d cytotoxicity in uveal melanoma cells, we compared our Cas13d-based approach with two other widely used knockdown and knockout methods in 92-1 cells: Cas9 nuclease-mediated gene knockout and siRNA-mediated knockdown. We introduced the Cas9 (*Streptococcus pyogenes*) knockout as a well-established all-in-one lentiviral transduction^50^, siRNA knockdown as transfecting a pool of RasGRP3-targeting siRNA^13^, and Cas13 as a combination of transduction (Cas13) and transfection (guide #4 or 3) similar to the approach in Figure 2D (**Fig. 4A**). Using live/dead staining followed by flow cytometry for quantification, we demonstrated that our Cas13d-based approach (viability reduced to <5%) outperformed both Cas9 knockout (∼45% viability) and siRNA knockdown (∼35% viability) (**Fig. 4B**). This data establishes that Cas13d–mediated knockdown elicits more potent cytotoxicity in RasGRP3-dependent UM cells than either DNA-level Cas9 knockout or siRNA-mediated knockdown.

**Figure 4:**
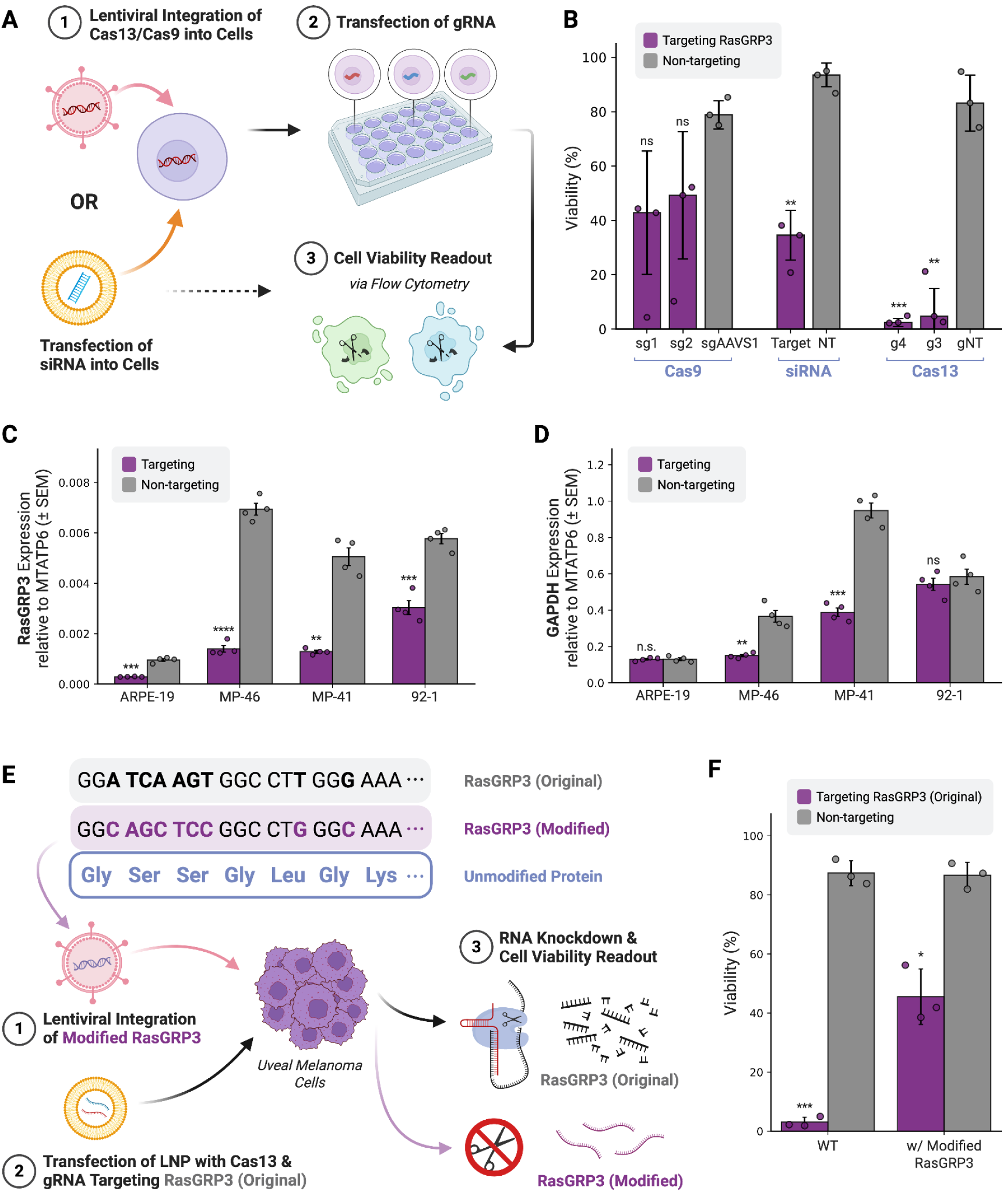
On-target mRNA knockdown and collateral cleavage by Cas13 contribute synergistically to cytotoxicity in cancer cells. (A) Schematic of methods for (B). Cas13d or SpCas9 are lentivirally integrated into 92-1 (UM) cells, followed by transfection of their corresponding guides. Alternatively, siRNA is transfected into the cells. Cell viability is quantified 48 hours after transfection by performing a Live/Dead stain followed by flow cytometry. (B) Comparison of Cas9 knockout, siRNA knockdown and Cas13d knockdown of RasGRP3 in 92-1 (UM) cells. Two RasGRP3-targeting guides and one non-targeting guide were tested for each CRISPR enzyme, along with RasGRP3-targeting siRNA and non-targeting siRNA. (C, D) qPCR results 12 hours after transfection of the Cas13d-LNP therapeutic into ARPE-19 (non-UM) and three UM cells (MP-46, MP-41, 92-1) (N=4). Expression relative to MTATP6 (mitochondrial RNA) is shown for (C) RasGRP3, and (D) GAPDH (off-target RNA for collateral cleavage). (E) Schematic of methods for (F). A sequence-modified, guide-orthogonal RasGRP3 gene (i.e., not targetable by targeting guide, but the same protein is produced) is stably integrated into uveal melanoma cells (92-1). The Cas13d-LNP therapeutic is then transfected, leading to degradation of the original RasGRP3 mRNA, but not the modified version. Viability of the cells is then quantified 48 hours after transfection to compare the impacts of collateral cleavage and RasGRP3 dependency. (F) In-vitro viability assay in modified uveal melanoma cells to determine the mechanism of cell killing by Cas13d (RasGRP3 knockdown vs. collateral cleavage).

With Cas13d outperforming RasGRP3 knockout/knockdown alone, we hypothesized that collateral cleavage yields the improved cytotoxicity. To quantify on-target and collateral RNA degradation, ARPE-19 (non-UM) and three UM lines (MP-46, MP-41, 92-1) were transfected with the Cas13d-LNP therapeutic, and then their gene expression was assayed via RT-qPCR at 12 hours (**Fig. 4C-D**).

A key feature of collateral RNA cleavage is its universal activity to all RNAs, including house-keeping genes commonly used as internal controls (e.g., *GAPDH*), thus making normalization challenging. To solve this problem, we used mitochondrial genes, which have been demonstrated as shielded from Cas13 collateral cleavage due to their subcellular compartmentation,^29^ as internal controls. Using *MTATP6* (a mitochondrial gene) as a normalization control, we demonstrated that *RASGRP3* mRNA was robustly knocked down in all four cell lines, confirming significant on-target RNA degradation (**Fig. 4C**). *GAPDH* levels were also significantly reduced in MP-46 and MP-41 cells, with a modest decrease in 92-1, but remained nearly unchanged in ARPE-19 (**Fig. 4D**). These results indicate that Cas13d exhibits substantial collateral cleavage preferentially in UM cells, likely as a result of high RasGRP3 expression.

To separate the cytotoxic contribution of RasGRP3 degradation from collateral cleavage, we engineered 92-1 cells to stably express a “guide-orthogonal” RasGRP3 cDNA containing synonymous mutations in the g4 binding site (**Fig. 4E**). Upon transfection of the Cas13d-LNP therapeutic, wild-type 92-1 cells were almost entirely eliminated (3.03 ± 1.70% viability), whereas orthogonal-RasGRP3 92-1 cells were partially rescued (45.55 ± 9.39% viability) (**Fig. 4F**). This partial rescue indicates that collateral cleavage likely contributes a viability reduction of around 2-fold in UM cells.

Altogether, we demonstrated that simple, on-target knockout (Cas9) and knockdown (siRNA) can reduce UM cells’ viability by approximately 50% (**Fig. 4B**), while collateral cleavage alone also contributes about 50% viability reduction (**Fig. 4F**). Although it is hard to cross compare between different methods of knockdown/knockout, the effects of on-target knockdown of RasGRP3 and collateral cleavage seem to work synergistically in the context of Cas13d-mediated UM killing, as we consistently observed an over 20-fold viability reduction across various conditions (**Fig. 2D, 3D, 4B, 4F**). Therefore, it is clear that the combination of the two mechanisms results in the Cas13d therapeutic’s improved cytotoxicity in UM.

## Discussion

Here, we establish a complete “target-to-therapy” workflow that begins with computational identification of a context-specific essential gene, proceeds through rapid guide screening, and culminates in a potential Cas13d cancer therapeutic delivered via LNP (**Fig. 5**). Using RasGRP3 as a proof of concept in uveal melanoma, we show that a single gRNA (g4) co-delivered with Cas13d mRNA eliminates >97% of three independent UM cell lines while sparing non-UM cells and non-cancerous eye (RPE) cells. This Cas13d-LNP therapeutic exhibits superior potency relative to Cas9 knockout and siRNA knockdown, illustrating the unique capacity of Cas13d to transiently “drug” intracellular targets while inducing selective cytotoxicity by combining on-target knockdown with collateral RNA degradation.

**Figure 5:**
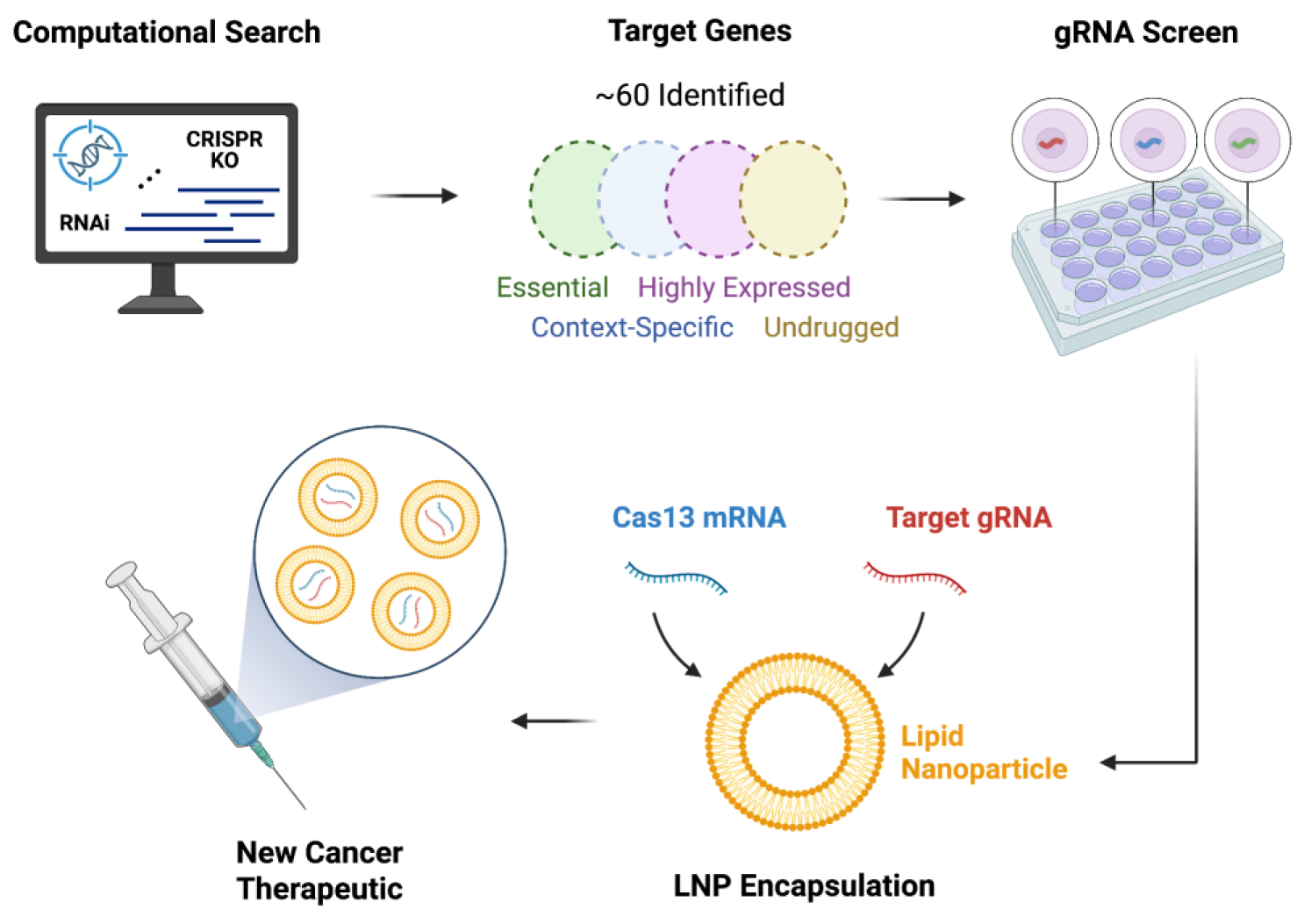
A pipeline for developing Cas13 therapeutics against cancer. A search through CRISPR-Cas9 KO, RNAi, and other cancer dependency screens yields initial targets, which can be filtered to find context-specific and undrugged fitness genes. Once the target has been identified, perform a gRNA screen to find the most effective/selective targeting guide. Finally, encapsulate the Cas13 mRNA and targeting gRNA into an LNP (or other delivery vehicle), creating a new Cas13 cancer therapeutic.

A four-phase pipeline for rapidly developing Cas13d-based cancer therapeutics is shown in **Figure 5**. First, large-scale dependency datasets—CRISPR–Cas9 knockouts, RNAi screens, and related functional genomics studies—are mined to compile candidate fitness genes, which are then filtered for cancer essentiality, context-specific lethality, high relative expression, and an absence of existing inhibitors/drugs. Next, a comprehensive library of Cas13d gRNAs are tiled across each transcript and screened in parallel viability assays to pinpoint gRNAs that maximize killing in target cells while sparing non-dependent lines. In the third phase, the lead gRNA is co-encapsulated with Cas13d mRNA into LNP formulations, and dozens of LNP compositions are evaluated for mRNA delivery efficiency and biocompatibility to select an optimized vehicle. Finally, the optimized LNP-Cas13d mRNA is tested across panels of cancerous and non-cancerous cell lines to confirm efficacy and specificity—laying the groundwork for *in vivo* validation. Due to our pipelines’ modularity—target identification, guide-RNA design, and LNP formulation—we expect it to be able to address a wide range of context-specific essential genes revealed by DepMap or comparable screens. Theoretically, a new cancer therapeutic could be created by simply changing the guide sequence.

Our results showed that LNP 3060i10 achieved ∼90% transfection efficiency in two UM cell lines without significant basal toxicity. Notably, delivery via LNP provides a clinically validated, redosing-compatible vehicle already deployed in mRNA vaccines and emerging CRISPR trials.^51,52^ Liver tropism of many clinically-validated LNPs might also be advantageous for UM, whose metastases localize to the liver in >90% of patients, yet direct ocular administration (i.e., suprachoroidal) remains attractive for early-stage disease.^53–55^ Future work should therefore evaluate two major features: (i) biodistribution and pharmacokinetics of the LNP-Cas13d mRNA in orthotopic ocular and hepatic xenograft models; and (ii) immune activation profiles after single and repeat dosing, particularly activation of innate RNA-sensing pathways that could curtail efficacy or cause inflammation.^56,57^

Rescue experiments with a guide-orthogonal RasGRP3 cDNA indicate that on-target knockdown and collateral cleavage contribute synergistically to cytotoxicity (**Fig. 4F**). Importantly, collateral activity was negligible in ARPE-19 cells, suggesting that the therapeutic window is widened by the high baseline expression of RasGRP3 in UM but not in healthy ocular tissue. This data suggests a two-threshold model. In non-cancerous cells, baseline RasGRP3 transcript levels remain below the activation threshold required to trigger robust Cas13d collateral cleavage; this limits Cas13d activity to the intended knockdown of a non-essential transcript and, thus, offers a self-limiting safety mechanism for normal tissues. In UM cells, high RasGRP3 expression pushes the system past this threshold, unleashing widespread RNA degradation that synergizes with direct target ablation to collapse essential signaling pathways.

There are a few limitations in the work. First, though we showed high efficacy data in all of three tested UM cell-line models, it is unknown whether our approaches can achieve similar efficacies in more complex models like *ex vivo* tumor organoids with heterogeneous RasGRP3 expression and LNP uptake. Second, guide-induced sequence variants escaping Cas13 recognition could emerge under selective pressure. A better strategy is to leverage the multiplexity of Cas13d or employ multiple gRNA cocktails to prevent resistance. Finally, we also expect that better efficacy can be achieved by combining our Cas13d therapeutic with other conventional therapies like immune checkpoint blockades or targeted kinase inhibitors.

To summarize, our work here demonstrates the power of coupling large-scale cancer dependency genomics/transcriptomics computational analysis with programmable RNA nucleases. Coupled with clinically validated lipid nanoparticles, we were able to transform previously “undruggable” intracellular proteins into tractable therapeutic targets. Unlike traditional drug development approaches that rely on small molecules or antibodies, the Cas13 system integrates seamlessly with computational pipelines—directly translating computationally identified targets into therapeutics, regardless of their druggability. By targeting RasGRP3 with Cas13d, we achieved rapid and highly-specific elimination of uveal melanoma cells, uncovered a dual-mechanism of action that exploits tumor-specific transcript abundance, and provided a generalizable roadmap for developing Cas13-based cancer therapeutics. Successful translation of this strategy to *in vivo* models—and ultimately to the clinic—could not only address the decades-long therapeutic void in uveal melanoma but also inaugurate a new class of RNA-targeting precision CRISPR therapeutics for cancer.

## Methods

### Cell Culture

Three UM cell lines—92-1 (CVCL_8607), MP-46 (CVCL_4D13), and MP-41 (CVCL_4D12)— were cultured in RPMI-1640 supplemented with 10% FBS, 1% penicillin–streptomycin, and 2 mM L-glutamine. Three non-UM cell lines—HEK 293T (CVCL_0063), HeLa (CVCL_0030), and ARPE-19 (CVCL_0145)—were cultured in DMEM/F-12 supplemented with 10% FBS, 1% penicillin–streptomycin, and 2 mM L-glutamine. All cells were cultured at 37 °C in 5% CO_2_. All accession codes are from Cellosaurus.

### Computational Identification of RasGRP3

Genome-wide Chronos dependency scores (DepMap 23Q4) were analyzed in Python 3.11. Genes with a Chronos score < −0.6 in at least one cell line were retained. Context-specific essential genes were called when any two of three tests—Welch’s t, Fisher’s exact (binary lethal/non-lethal) and Wilcoxon rank-sum—were significant (FDR < 0.05). Expression filters were manually applied based on GTEx and TCGA pan-cancer TPM data (≥ 2-fold over both healthy tissue and non-UM tumours). Based on data from Open Targets, previously drugged or easily druggable genes were excluded.

### Cas13d Guide (gRNA) Design

30-mer gRNAs tiled across RasGRP3 (ENSG00000152689) were generated using the Arc Institute’s “Deep Learning-Powered Cas13d Guide Design” web app and NYGC’s Cas13design. 42 of the top-ranked guides were chosen in order to target variable regions of the transcript. These guide spacers were ordered as oligonucleotides (IDT) and cloned into a pU6-RfxCas13d-crRNA plasmid via BsmBI (NEB); all inserts were sequence-verified.

### Lentiviral Transduction of Cas13d

The RfxCas13d-P2A-mCherry cassette was cloned into pLenti-EF1α. To produce lentivirus, Lenti-X cells were seeded in 6-well plates and cultured overnight in 2 mL DMEM medium (defined above). 850 μl culture medium was removed and cells were transfected with 0.55 μg pMD2.G (Addgene #12259), 1.28 μg psPAX2 (Addgene #12260), and 1.79 μg Cas13d plasmid in 426 μl Opti-MEM (Gibco) using 10.88 μl TransIT-LT1 (Mirus Bio). Six hours later, the culture medium was completely removed and replaced with fresh DMEM medium and 1x ViralBoost (Alstem Bio). At 24 hours post-transfection, viral supernatant was harvested and filtered through a 0.45 μm syringe filter. The viral supernatant was mixed with Lentivirus Precipitation Solution (Alstem Bio), incubated at 4°C for at least 4 hours, and concentrated 10x in Opti-MEM. Concentrated virus was either used fresh or kept frozen at −80 °C for up to 48 hours. UM and control cells seeded in 24-well plates were transduced with 1/3 of the produced lentivirus per well. Viral media was replaced after 48 hours, yielding stably-integrated Cas13d cell lines.

### Guide RNA Transfection and Viability Assay

Cas13d-transduced cells were seeded in 96-well plates (0.1 × 10^5^ per well) and transfected with 100 ng gRNA plasmid using Mirus TransIT-LT1. At 96 hours, cell viability was quantified using LIVE/DEAD™ Fixable Far Red Dead Cell Stain Kit (ThermoFisher), followed by flow cytometry. Gating of live/dead cells was defined by wild-type controls with ∼100% live cells that were stained with the same reagent mentioned above.

### LNP Manufacturing

Cas13d mRNA was synthesized via in vitro transcription (IVT) using the HiScribe T7 mRNA Kit with CleanCap Reagent AG (NEB). LNP 3060i10 was formulated by ethanol injection, combining an organic phase composed of 48.45% 306O_i10_, 19.43% DOPE, 31.16% cholesterol, and 0.96% DMG-PEG2000, with an aqueous phase containing Cas13d mRNA in 25 mM acetate buffer, at a 1:3 volume ratio and a 16:1 weight ratio. Cells were treated by directly adding the mRNA/LNP formulation to the culture medium at a final concentration of 10 µg per million cells. LNP library screening was conducted in a similar manner, with variations in the ionizable lipid component of the organic phase. Transfection efficiency was assessed based on GFP mRNA expression.

### LNP Transfection Efficiency

GFP mRNA-loaded LNPs (10 µg per million cells) were added to target cells; GFP+ percentages were quantified at 48 hours by flow cytometry. 30 LNP formulations with varying ionizable lipid chemotypes were screened.

### Cas13d-LNP Viability Assays

For single-dose cytotoxicity, cells were treated with Cas13d/g4 LNPs (as detailed above). At 48 hours, viability was assessed by the Live/Dead assay (detailed above). For collateral-cleavage studies, total RNA was extracted at 12 hours for RT-qPCR.

### Cas9 and siRNA Benchmarking

For DNA knockout, 92-1 cells were lentivirally transduced (method explained above) with one of three plasmids, each containing spCas9 and one of three sgRNAs (two different targeting guides and one non-targeting guide) constitutively expressed under separate promoters (**Supp. Table S3**). Viability was measured 96 hours after transduction. siRNA pools (Dharmacon, RasGRP3-targeting and NT SMARTpools) were transfected with Lipofectamine RNAiMAX (30 nM). Viability was measured 48 hours after transfection.

### Gene Expression Quantification with RT-qPCR

For quantification of gene expression via RT-qPCR, cells were harvested at the indicated time point and extracted total RNA (Zymo cat#R1055), then reverse transcribed (NEB cat#E3010) them with a poly-dT primer (IDT cat#51-01-15-08). The qPCR reactions were performed with a standard SYBR Green system (Bio-Rad cat#1725270). One pair of primers was used for each of GAPDH, MTATP6, and MTND1, while two primer pairs were used (separately) for RasGRP3. The specific primers used in this study are provided in **Supp. Table S4**.

### Orthogonal RasGRP3 for Collateral Cleavage Study

A codon-optimized RasGRP3 cDNA (VectorBuilder Codon Optimization Tool) was generated and manually examined to ensure sufficient mutations to lack complementarity with the g4 gRNA spacer. The modified RasGRP3 was cloned into a pEF1α-GFP-Puro backbone. 92-1 cells were transduced and then selected in 1 µg mL^-1^ puromycin for seven days. Rescue and wild-type 92-1 cells were then treated with the Cas13d/g4 LNP (3060i10; 1 µg RNA well^-1^, 12-well plate), and viability was measured 48 hours later.

### Statistical Analysis and Quantification

Unless otherwise stated, data are mean ± SEM from N=3 biologically independent replicates. Statistical significance between groups was determined using two-tailed unpaired Welch’s t-tests and was conducted using SciPy v1.15.3. Unless otherwise mentioned, statistical tests were performed between a targeting condition and its non-targeting (or other negative) control. When applicable, p-values were adjusted for multiple comparisons.

All plots were generated in Python 3.11. Bar graphs show the mean with error bars representing the standard error of the mean (SEM). Box plots were generated using ggplot2 and indicate the median value, along with the first and third quartiles (IQR). Whiskers are limited to 1.5 times the distance of the IQR. In figures, asterisks are used to represent the following significance thresholds: p < 0.05 (*), < 0.01 (**), < 0.001 (***), < 0.0001 (****).

## Data and Code Availability

The datasets used in this study to identify RasGRP3 and other context-specific essential genes are provided by the Broad Institute’s DepMap. RNA Expression datasets can be obtained online from TCGA and GTEx. Analysis details are provided in the Methods section. Requests for additional data and resources should be directed to and will be fulfilled by the lead contact, Lei S. Qi.

## Acknowledgements

The authors thank Dr. Fabrizia Urbinati, Dr. Mark Smith, Dr. Vinit Mahajan, and Dr. Prithvi Mruthrunjaya for helpful discussions and advice. The authors thank Dr. Vinit Mahajan for providing uveal melanoma and RPE cell lines. The authors thank all members of the Lei Stanley Qi lab for helpful discussions, especially T. Wachsmann, G. Roth, J. Magnusson, and J. Bezney. D.S. and L.S. thank the ChEM-H Undergraduate Entrepreneurship Program for support. Y.M. is supported by a Cancer Research Institute Dr. Keith Landesman Memorial Fellowship (CRI5247). D.M. acknowledges support from departmental core grants from the NIH (P30-EY026877) and an unrestricted grant from Research to Prevent Blindness. L.S.Q. acknowledges support by National Science Foundation (NSF), National Institutes of Health (NIH), and Chau Hoi Shuen Foundation. Research was supported by the National Institute of Neurological Disorders and Stroke of NIH (DP1NS137219), National Cancer Institute (R01CA266470), and NSF CAREER award (2046650). L.S.Q. is a Chan Zuckerberg Biohub investigator.

## Author Contributions

D.S. conceived of the original idea. D.S. and L.S. contributed significantly to the final design with input from D.M. and L.S.Q. D.S., L.S., and Y.M. contributed significantly to the experimental plan. D.S. performed all computational work, cloned plasmids, collected and analyzed all data, performed all work with cell cultures, produced lentivirus, and performed most experiments. L.S. helped clone gRNAs and perform the guide screen. Y.M. cloned the Cas9 plasmids and produced lentivirus. S.P. and A.L. cloned simplified gRNAs and Cas13d, performed IVT, formulated LNPs, and encapsulated mRNA in LNPs. D.S. and L.S. created the figures and secured funding for the project. L.S.Q. supervised the work. D.S. and L.S. wrote the manuscript, and L.S.Q. revised it with edits from Y.M. and S.P. and input from all authors.

## Declaration of interests

D.S. and L.S.Q. are inventors of a provisional patent that is being filed based on this work via Stanford University. L.S.Q. is a founder of Epicrispr Biotechnologies and scientific advisor of Laboratory of Genomic Research.

## Supplementary figures and captions

**Figure S1:**
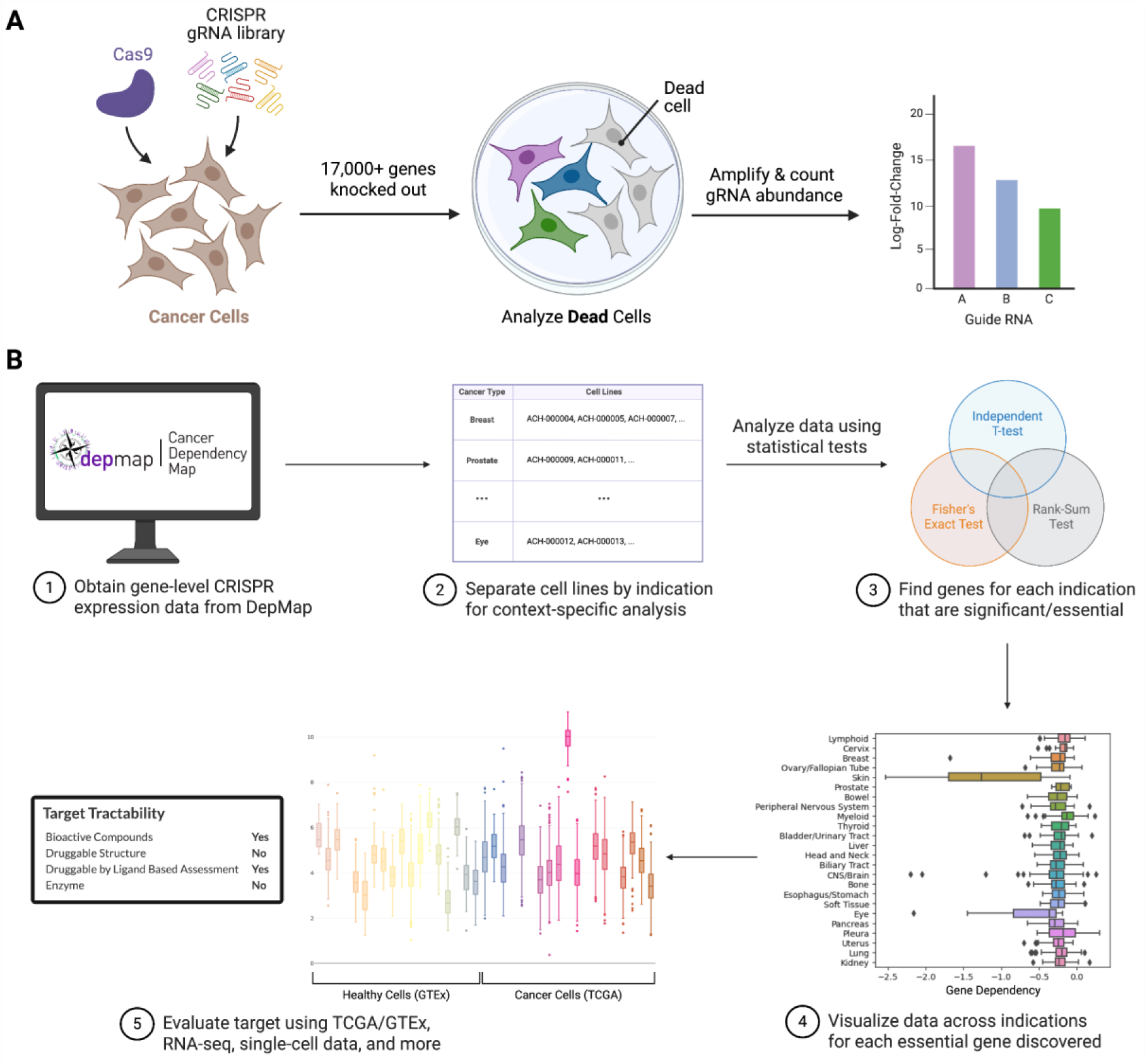
Statistical analysis of CRISPR screen datasets to uncover context-specific essential cancer genes. (**A**) Schematic of methods used in large-scale CRISPR-Cas9 KO screens like DepMap. The final gRNA abundance values are used to determine Chronos (dependency) scores. (**B**) Computational workflow to identify context-specific essential genes for cancer, followed by further investigation into their therapeutic potential and druggability. Data is initially obtained from cancer dependency screens like DepMap. Cell lines are then grouped into cancer indications to run statistical tests between indications. Significance in at least two out of three different statistical tests designates the gene as a context-specific essential gene. Finally, the therapeutic potential and druggability of the gene can be further validated using RNA-seq datasets, target databases, and more.

**Figure S2:**
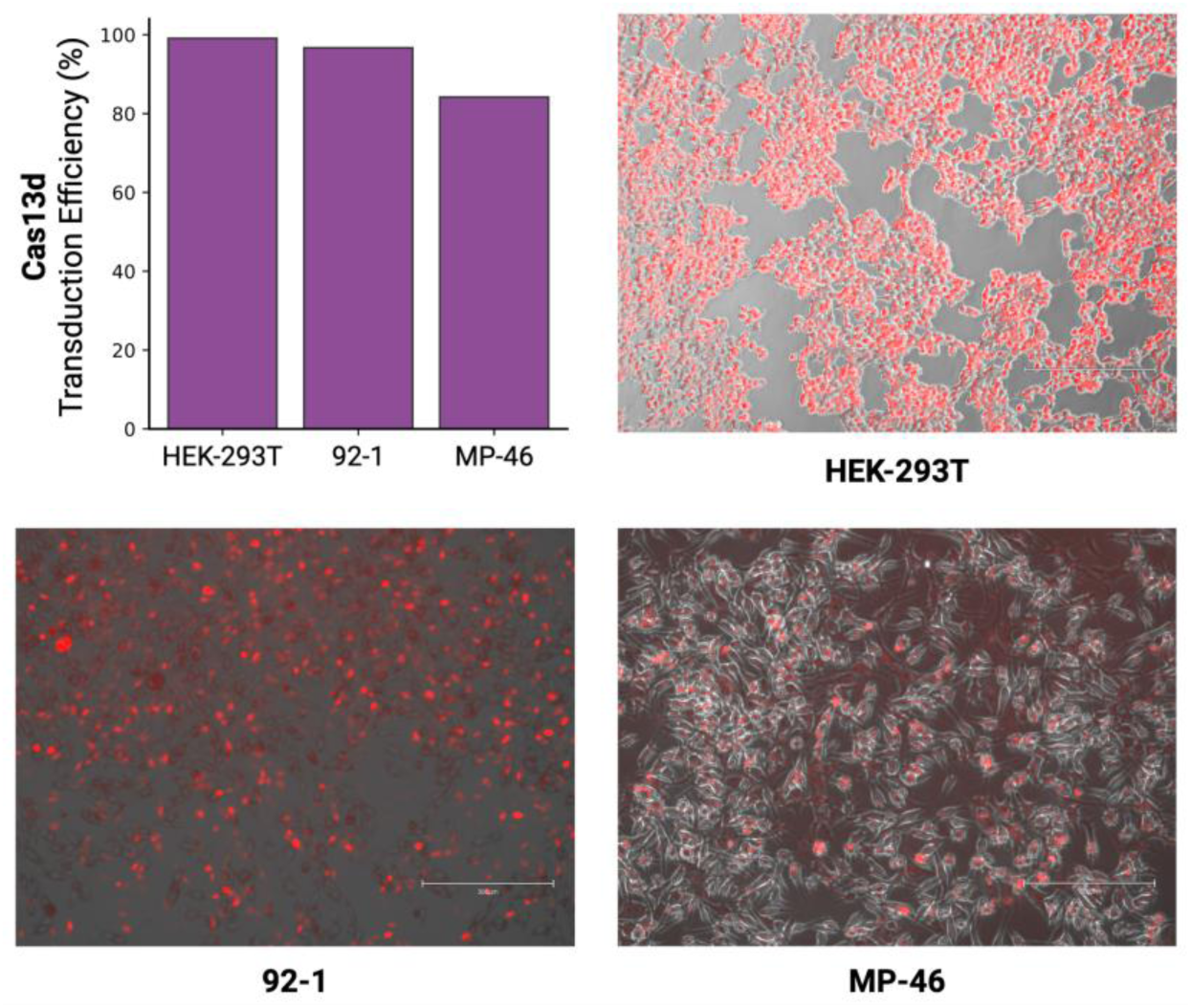
Lentiviral transduction efficiency of Cas13d in uveal melanoma cells and HEK-293T cells. A Cas13d-P2A-mCherry construct is lentivirally integrated into HEK-293T, 92-1, and MP-46 cells. Transduction efficiency is quantified by flow cytometry, with a gate for mCherry+ cells defined by untransduced controls.

**Figure S3:**
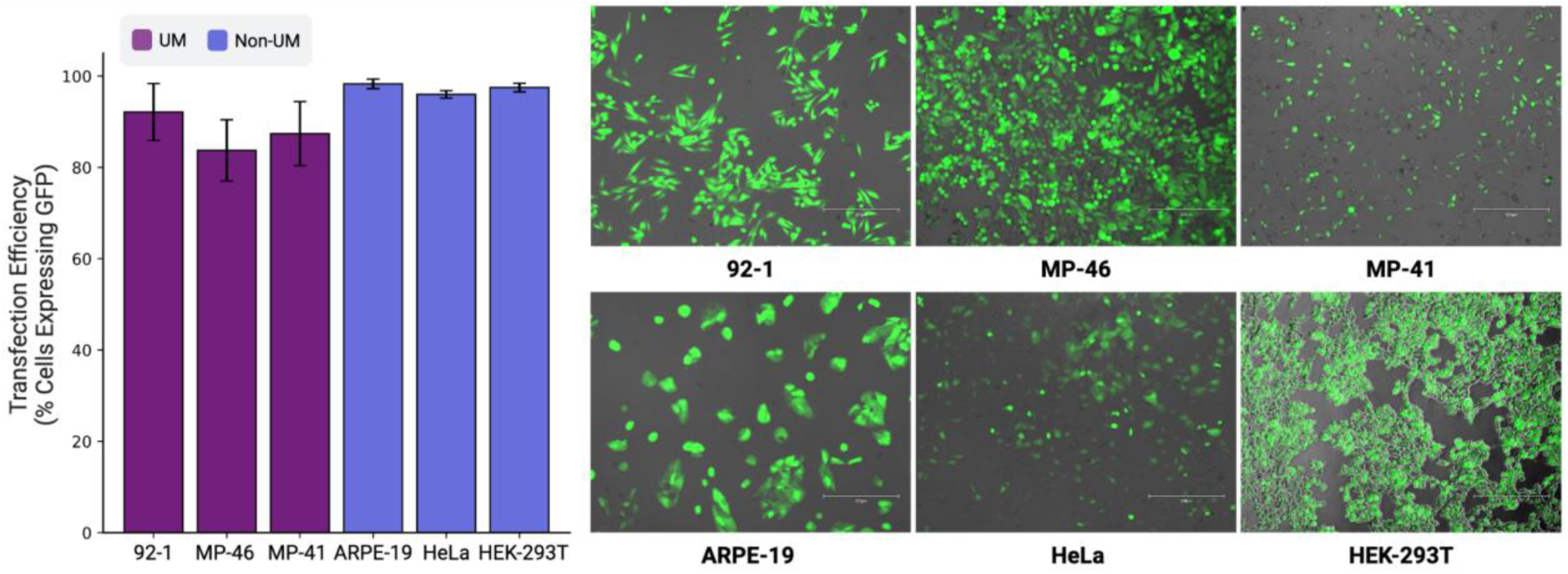
Transfection efficiency of the top LNP candidate (3060i10) in UM and non-UM cell lines. GFP mRNA is transfected into three UM cell lines (92-1, MP-46, MP-41) and three non-UM cell lines (ARPE-19, HeLa, HEK-293T). Transfection efficiency is quantified by flow cytometry, with a gate for GFP+ cells defined by untransfected controls.

**Figure S4:**
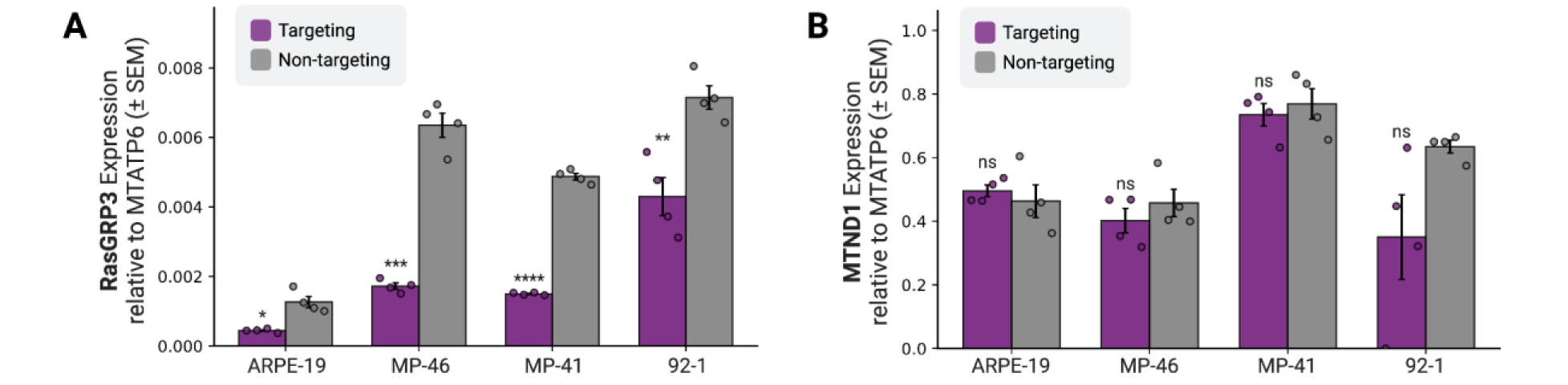
Additional RT-qPCR results after transfection of the Cas13d-LNP. The therapeutic was transfected into ARPE-19 (non-UM) and three UM cells (MP-46, MP-41, 92-1). Expression relative to MTATP6 (mitochondrial RNA) is shown for **(A)** RasGRP3 (using a second set of primers), and **(B)** MTND1 (another mitochondrial RNA, which should match up with MTATP6).

## Notes

### Competing Interest Statement

D.S. and L.S.Q. are inventors of a provisional patent related to this work via Stanford University. L.S.Q. is a founder of Epicrispr Biotechnologies and scientific advisor of Laboratory of Genomic Research.

## References

1. Jager, M. J. et al. Uveal melanoma. Nature Reviews Disease Primers 6, 1–25 (2020).

2. Patel, D. R., Blair, K. & Patel, B. C. Ocular Melanoma. in StatPearls [Internet] (StatPearls Publishing, 2024).

3. Kaliki, S., Shields, C. L. & Shields, J. A. Uveal melanoma: estimating prognosis. Indian J Ophthalmol 63, 93–102 (2015).

4. Singh, A. D., Turell, M. E. & Topham, A. K. Uveal melanoma: trends in incidence, treatment, and survival. Ophthalmology 118, (2011).

5. Roelofsen, C. D. M. et al. Five Decades of Enucleations for Uveal Melanoma in One Center: More Tumors with High Risk Factors, No Improvement in Survival over Time. Ocul Oncol Pathol 7, 133–141 (2021).

6. Lane, A. M., Kim, I. K. & Gragoudas, E. S. Survival Rates in Patients After Treatment for Metastasis From Uveal Melanoma. JAMA Ophthalmology 136, 981–986 (2018).

7. Branisteanu, D. C. et al. Uveal melanoma diagnosis and current treatment options (Review). Experimental and Therapeutic Medicine 22, 1428 (2021).

8. Hassel, J. C. et al. Three-Year Overall Survival with Tebentafusp in Metastatic Uveal Melanoma. N Engl J Med 389, 2256–2266 (2023).

9. Sacco, J. J. et al. Long-term survival follow-up for tebentafusp in previously treated metastatic uveal melanoma. Journal for Immunotherapy of Cancer 12, e009028 (2024).

10. Chen, L. N. & Carvajal, R. D. Tebentafusp for the treatment of HLA-A*02:01-positive adult patients with unresectable or metastatic uveal melanoma. Expert Rev Anticancer Ther 22, 1017–1027 (2022).

11. Saldanha, E. F. et al. How we treat patients with metastatic uveal melanoma. ESMO Open 10, 104496 (2025).

12. Kaliki, S. & Shields, C. L. Uveal melanoma: relatively rare but deadly cancer. Eye 31, 241– 257 (2016).

13. Chen, X. et al. RasGRP3 Mediates MAPK Pathway Activation in GNAQ Mutant Uveal Melanoma. Cancer cell 31, 685 (2017).

14. Silva-Rodríguez, P. et al. GNAQ and GNA11 Genes: A Comprehensive Review on Oncogenesis, Prognosis and Therapeutic Opportunities in Uveal Melanoma. Cancers 14, 3066 (2022).

15. Smit, K. N., Jager, M. J., de Klein, A. & Kiliҫ, E. Uveal melanoma: Towards a molecular understanding. Progress in Retinal and Eye Research 75, 100800 (2020).

16. Tsherniak, A. et al. Defining a Cancer Dependency Map. Cell 170, 564–576.e16 (2017).

17. Arafeh, R., Shibue, T., Dempster, J. M., Hahn, W. C. & Vazquez, F. The present and future of the Cancer Dependency Map. Nature Reviews Cancer 25, 59–73 (2024).

18. Hart, T. et al. High-Resolution CRISPR Screens Reveal Fitness Genes and Genotype-Specific Cancer Liabilities. Cell 163, 1515–1526 (2015).

19. Behan, F. M. et al. Prioritization of cancer therapeutic targets using CRISPR–Cas9 screens. Nature 568, 511–516 (2019).

20. Pacini, C. et al. A comprehensive clinically informed map of dependencies in cancer cells and framework for target prioritization. Cancer Cell 42, 301–316.e9 (2024).

21. Dang, C. V., Reddy, E. P., Shokat, K. M. & Soucek, L. Drugging the ‘undruggable’ cancer targets. Nature Reviews Cancer 17, 502–508 (2017).

22. Lu, Y. et al. Emerging Pharmacotherapeutic Strategies to Overcome Undruggable Proteins in Cancer. Int J Biol Sci 19, 3360–3382 (2023).

23. Moore, A. R. et al. GNA11 Q209L Mouse Model Reveals RasGRP3 as an Essential Signaling Node in Uveal Melanoma. Cell Reports 22, 2455–2468 (2018).

24. Abudayyeh, O. O. et al. RNA targeting with CRISPR–Cas13. Nature 550, 280–284 (2017).

25. Konermann, S. et al. Transcriptome Engineering with RNA-Targeting Type VI-D CRISPR Effectors. Cell 173, 665–676.e14 (2018).

26. Zhu, G. et al. CRISPR–Cas13: Pioneering RNA Editing for Nucleic Acid Therapeutics. Biodesign Research 6, 0041 (2024).

27. Du, S. W. & Palczewski, K. Eye on genome editing. Journal of Experimental Medicine 220, e20230146 (2023).

28. Murphy, R. & Martin, K. R. Genetic engineering and the eye. Eye 39, 57–68 (2024).

29. Shi, P. et al. Collateral activity of the CRISPR/RfxCas13d system in human cells. Communications Biology 6, 1–8 (2023).

30. Ai, Y., Liang, D. & Wilusz, J. E. CRISPR/Cas13 effectors have differing extents of off-target effects that limit their utility in eukaryotic cells. Nucleic Acids Res 50, e65 (2022).

31. Bot, J. F., van der Oost, J. & Geijsen, N. The double life of CRISPR–Cas13. Current Opinion in Biotechnology 78, 102789 (2022).

32. Mashel, T. V. et al. Overcoming the delivery problem for therapeutic genome editing: Current status and perspective of non-viral methods. Biomaterials 258, 120282 (2020).

33. Tenchov, R., Bird, R., Curtze, A. E. & Zhou, Q. Lipid Nanoparticles─From Liposomes to mRNA Vaccine Delivery, a Landscape of Research Diversity and Advancement. ACS Nano (2021) doi:10.1021/acsnano.1c04996.

34. Hou, X., Zaks, T., Langer, R. & Dong, Y. Lipid nanoparticles for mRNA delivery. Nature Reviews Materials 6, 1078–1094 (2021).

35. El-Mayta, R., Padilla, M. S., Billingsley, M. M., Han, X. & Mitchell, M. J. Testing the In Vitro and In Vivo Efficiency of mRNA-Lipid Nanoparticles Formulated by Microfluidic Mixing. J Vis Exp (2023) doi:10.3791/64810.

36. Wang, J. et al. Recent Advances in Lipid Nanoparticles and Their Safety Concerns for mRNA Delivery. Vaccines (Basel) 12, (2024).

37. Yan, J., Kang, D. D. & Dong, Y. Harnessing lipid nanoparticles for efficient CRISPR delivery. Biomaterials science 9, 6001 (2021).

38. Chambers, C. Z. et al. Lipid Nanoparticle-Mediated Delivery of mRNA Into the Mouse and Human Retina and Other Ocular Tissues. Transl Vis Sci Technol 13, 7 (2024).

39. Zabaleta, N., Torella, L., Weber, N. D. & Gonzalez-Aseguinolaza, G. mRNA and gene editing: Late breaking therapies in liver diseases. Hepatology (Baltimore, Md.) 76, 869 (2022).

40. DepMap: The Cancer Dependency Map Project at Broad Institute. https://depmap.org/portal/.

41. Weinstein, J. N. et al. The Cancer Genome Atlas Pan-Cancer analysis project. Nature Genetics 45, 1113–1120 (2013).

42. The Cancer Genome Atlas Program (TCGA). https://www.cancer.gov/ccg/research/genome-sequencing/tcga (2022).

43. Lonsdale, J. et al. The Genotype-Tissue Expression (GTEx) project. Nature Genetics 45, 580–585 (2013).

44. Buniello, A. et al. Open Targets Platform: facilitating therapeutic hypotheses building in drug discovery. Nucleic Acids Res 53, D1467–D1475 (2025).

45. RASGRP3 RAS guanyl releasing protein 3 [Homo sapiens (human)] - Gene - NCBI. https://www.ncbi.nlm.nih.gov/gene/25780.

46. Open Targets Platform. https://platform.opentargets.org/.

47. Wei, J. et al. Deep learning and CRISPR-Cas13d ortholog discovery for optimized RNA targeting. Cell Syst 14, 1087–1102.e13 (2023).

48. GitHub - Arc Institute Cas13d guide design. GitHub https://github.com/ArcInstitute/RNAtargeting_web_custom.

49. Wessels, H.-H. et al. Massively parallel Cas13 screens reveal principles for guide RNA design. Nature Biotechnology 38, 722–727 (2020).

50. Sanjana, N. E., Shalem, O. & Zhang, F. Improved vectors and genome-wide libraries for CRISPR screening. Nat Methods 11, 783–784 (2014).

51. Polack, F. P. et al. Safety and Efficacy of the BNT162b2 mRNA Covid-19 Vaccine. New England Journal of Medicine (2020) doi:10.1056/NEJMoa2034577.

52. Gillmore, J. D. et al. CRISPR-Cas9 In Vivo Gene Editing for Transthyretin Amyloidosis. The New England journal of medicine 385, (2021).

53. Eygeris, Y., Gupta, M., Kim, J. & Sahay, G. Chemistry of Lipid Nanoparticles for RNA Delivery. Accounts of Chemical Research (2021) doi:10.1021/acs.accounts.1c00544.

54. Woodman, S. E. Metastatic uveal melanoma: biology and emerging treatments. Cancer J 18, 148–152 (2012).

55. Uveal melanoma: In the era of new treatments. Cancer Treatment Reviews 119, 102599 (2023).

56. Uehata, T. & Takeuchi, O. RNA Recognition and Immunity-Innate Immune Sensing and Its Posttranscriptional Regulation Mechanisms. Cells 9, (2020).

57. Kong, L.-Z. et al. Understanding nucleic acid sensing and its therapeutic applications. Experimental & Molecular Medicine 55, 2320–2331 (2023).

